# Endogenous suppression of WNT signalling in human embryonic stem cells leads to low differentiation propensity towards definitive endoderm

**DOI:** 10.1101/2020.07.03.186932

**Authors:** Dominika Dziedzicka, Mukul Tewary, Alexander Keller, Laurentijn Tilleman, Laura Prochazka, Joel Östblom, Edouard Couvreu De Deckersberg, Christina Markouli, Silvie Franck, Filip Van Nieuwerburgh, Claudia Spits, Peter W. Zandstra, Karen Sermon, Mieke Geens

## Abstract

Low differentiation propensity towards a targeted lineage can significantly hamper the utility of individual human pluripotent stem cell (hPSC) lines in biomedical applications. Here, we use monolayer and micropatterned cell cultures, as well as transcriptomic profiling, to investigate how variability in signalling pathway activity between human embryonic stem cell lines affects their differentiation efficiency towards definitive endoderm (DE). We show that endogenous suppression of WNT signalling in hPSCs at the onset of differentiation prevents the switch from self-renewal to DE specification. Gene expression profiling reveals that this inefficient switch is reflected in *NANOG* expression dynamics. Importantly, we demonstrate that higher WNT stimulation or inhibition of the PI3K/AKT signalling can overcome the DE commitment blockage. Our findings highlight that redirection of the activity of Activin/NODAL pathway by WNT signalling towards mediating DE fate specification is a vulnerable spot, as disruption of this process can result in poor hPSC specification towards DE.

## Introduction

The use of human pluripotent stem cells (hPSCs) in biomedical applications is hampered by variable efficiencies with which individual lines differentiate towards desired cell lineages (Bock et al., 2011; Hu et al., 2010; Kim et al., 2007; Osafune et al., 2008). Both (epi)genetic and environmental factors can contribute to this functional variability (Keller et al., 2018; Ortmann and Vallier, 2017). For example, differences in genetic, epigenetic and transcriptomic profiles between individual hPSC lines (Adewumi et al., 2007; Bock et al., 2011; Kilpinen et al., 2017; Markouli et al., 2019; Skottman et al., 2005) can lead to different levels of activity of signalling pathways, resulting in an individual line’s differential response to differentiation cues.

Human PSC differentiation efficiency can be improved by optimising differentiation conditions for individual lines (Hu et al., 2010; Kattman et al., 2011), or by screening and selecting lines with the highest differentiation efficiency for an intended application. While specific expression profiles at the undifferentiated stage can act as indicators of hPSC differentiation propensity (Jiang et al., 2013; Kim et al., 2011; Ran et al., 2013), not all disruptions to differentiation programmes are detectable at this stage. Therefore, a more optimal screening approach can be to evaluate lines based on their early lineage specification efficiency. Moreover, studying mechanisms which lead to differentiation bias can improve our knowledge about key signalling pathways involved in hPSC fate specification and lead to further improvement of differentiation protocols.

Given the importance of querying the differentiation potential of multiple hPSC lines, tools have been developed to address this question in a standardized manner. Examples include the TeratoScore, which quantitatively assesses trilineage differentiation capacity based on gene expression signatures of hPSC-derived teratomas (Avior et al., 2015), and the ScoreCard, which scores the gene expression profiles of differentiating embryoid bodies (EBs; Bock et al., 2011; Tsankov et al., 2015). However, both methods are suboptimal for high-throughput studies because they are time-consuming, taking from 7 days (ScoreCard) to several weeks (TeratoScore). In contrast, a recently developed *in vitro* platform allows for the generation of peri-gastrulation-like fate patterning in geometrically-confined colonies within only 2 days (Tewary et al., 2017, 2018), and has been recently validated for robust and quantitative screening of differentiation propensities of multiple hPSC lines (Tewary et al., 2019). Thus, this micropatterned differentiation can be a valuable alternative to the previously established tools.

Efficient differentiation of hPSCs towards endodermal lineages, e.g. of liver or pancreas, is of great value to regenerative medicine and drug development. Most established differentiation protocols initiate definitive endoderm (DE) specification by modulating the activity of Activin/NODAL and WNT signalling (D’Amour et al., 2006; Pagliuca et al., 2014; Rezania et al., 2012), though some additionally employ modifiers of FGF, BMP and/or PI3K/AKT pathway activity (Hannan et al., 2013; Loh et al., 2014). The Activin/NODAL pathway is known to play an important role in both maintaining pluripotency and guiding the cells towards DE specification (Bertero et al., 2015; Brown et al., 2011), whereas the role of WNT signalling in DE differentiation is to switch the specificity of Activin/NODAL signalling from supporting pluripotency to initiating mesendoderm specification (D’Amour et al., 2006; Funa et al., 2015; Yoney et al., 2018).

Our study aimed to uncover how variability in the activity of key signalling pathways associated with either maintenance of pluripotency or early lineage specification influences the differentiation propensity of individual hPSC lines towards DE. We used short-term monolayer differentiation protocols and the peri-gastrulation-like patterning to classify our human embryonic stem cell (hESC) lines according to their early differentiation propensity towards DE lineages and subsequently evaluated their transcriptomic profiles. Our results show that endogenous suppression of WNT signalling can result in reduced hPSC differentiation propensity towards DE and highlights that the essential switch of Activin/NODAL activity is a vulnerable spot on the way to efficient DE specification.

## Results

### Differentiation propensity screen identifies a hESC line with poor differentiation efficiency towards DE

We first screened a panel of four hESC lines (VUB01, VUB02, VUB03 and VUB04), routinely grown on human recombinant laminin-521 (LN521), for their efficiency to differentiate towards DE. We used established directed differentiation protocols towards mesendoderm, DE and hepatic progenitors (Cameron et al., 2015; D’Amour et al., 2005; Sui et al., 2012) adapted to LN521-based culture (**Fig. 1A**). After the first 24h of mesendoderm differentiation, we detected differences in the induction of the primitive streak marker *Brachyury (T*), with VUB03 and VUB04 lines showing the lowest but variable upregulation (**Fig. 1B, C**). In the subsequent 48h differentiation towards DE, VUB04 performed the poorest (**Fig. 1D-F, Fig. S1A**), with significantly lower expression of DE markers *SOX17* (on average 6.25-fold decrease) and *FOXA2* (on average 4.03-fold decrease) in comparison to the other three lines. VUB04 also retained high expression (6.11-fold increase) of the pluripotency marker *POU5F1* (**Fig. 1D**). Consistently, at the protein level, less than 10% of the VUB04 cells were positive for SOX17, whereas other lines showed a range of 65-80% SOX17-positive cells (**Fig. 1E**). In addition, 85% of VUB04 cells still expressed POU5F1, which was significantly higher than the other lines, suggesting that VUB04 did not efficiently exit the pluripotent state. Next, we subjected VUB04 to 8-day hepatic progenitor (HP) differentiation to establish whether the inability to specify towards DE lineages hampered the specification of DE derivatives. As suspected, expression of hepatic progenitor markers *HNF4*α, *AFP* and *SOX17* was significantly lower and the expression of pluripotency markers *POU5F1* and *SOX2* significantly higher in VUB04 in comparison to VUB01 (**Fig. 1G, H**).

**Figure 1.**
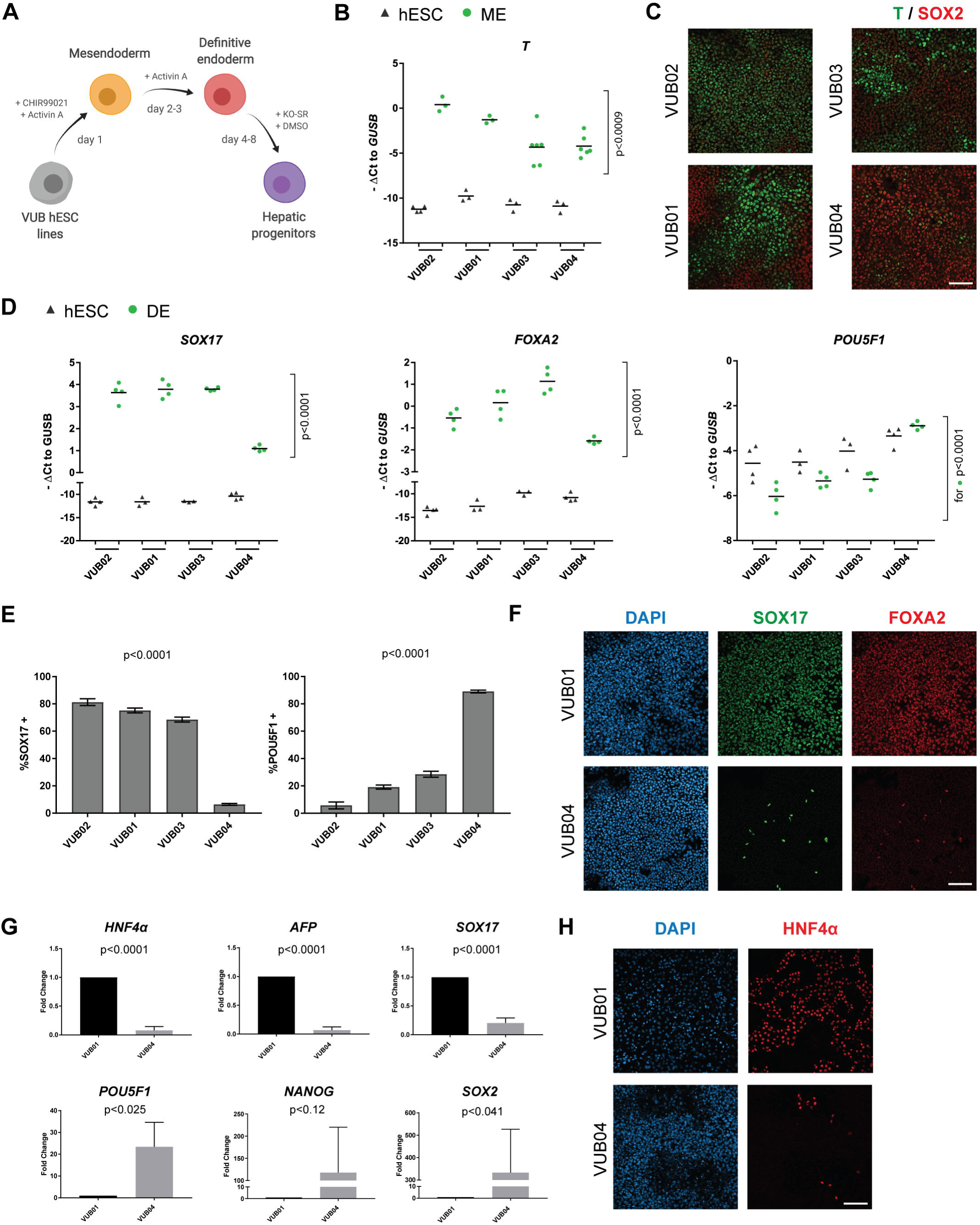
Differentiation propensity screen of four hESC lines shows that the VUB04 line has a low differentiation efficiency towards definitive endoderm (DE). A) Schematic overview of differentiation protocols used to direct the cells towards mesendoderm (ME), DE and hepatic progenitors (HP). Created with BioRender. B) Gene expression levels of *T* in hESCs and ME samples. Black triangles and green dots represent undifferentiated and differentiated samples, respectively. C) Representative immunofluorescent images for T and SOX2 after 24-hour ME differentiation. D) Expression level of *SOX17, FOXA2* and *POU5F1* in hESCs and DE samples. Black triangles and green dots represent undifferentiated and differentiated samples, respectively. E) Percentage of SOX17- and POU5F1-positive cells after 72-hour DE differentiation of hESC lines. F) Representative immunofluorescent images for SOX17 and FOXA2 after DE differentiation of VUB01 and VUB04. G) Comparison of expression levels of hepatic, DE and pluripotency markers between VUB01 and VUB04 in HP samples. H) Representative immunostainings for HNF4α after 8-day HP differentiation of VUB01 and VUB04. All scale bars represent 100 μm. All gene expression data and the quantified immunofluorescent data are representative of at least three (panel B and D) or three (panel G) biological replicates. The p-values were calculated using either one-way ANOVA (panel B, D and E) or unpaired t-test (panel G). For panels B and D, the p-values were calculated only for differentiated samples (green dots).

To check whether the differentiation impairment of VUB04 is specific towards the DE lineage or more general, we differentiated all four hESC lines to the neuroectoderm lineage (**Fig. S1B**). Although some variability in the expression levels of *PAX6* and *SOX1* was observed, VUB04 did not differ significantly from the other lines (**Fig. S1C, D**). Additionally, we performed 12-day spontaneous EB differentiation in serum-free APEL™ medium, followed by gene expression profiling with the ScoreCard assay. For this, we generated equal-sized aggregates following our previously published protocol (Dziedzicka et al., 2016) to avoid any potential differentiation bias originating from differentially sized EBs. Intriguingly, the ScoreCard indicated that all VUB lines, including VUB04, significantly downregulated pluripotency genes and upregulated genes associated with the three germ layers (**Fig. S1E**). As in this experiment the lines were subjected to spontaneous differentiation in a 3-dimensional environment, the result indicated that VUB04 does not efficiently differentiate towards DE upon exposure to defined modulators of signalling pathways in conventional monolayer cultures.

### Micropatterned differentiation confirms hESC differentiation propensity

To confirm that VUB04 has an impaired response to DE specification cues, we subjected our lines to directed differentiation in an alternative culture system. We used a micropatterning technology which allows for precise control of spatial microenvironment by confining cells to defined circular geometries. It has been recently demonstrated that in this system, upon BMP4 induction, micropatterned circular colonies of hPSCs form radially segregated regions of a population of cells retaining SOX2 expression in the centre and a ring of T-positive cells close to the edge (Tewary et al., 2017). Additionally, we recently showed that by modifying culture conditions it is also possible to obtain the DE population within these micropatterns, with cells double positive for SOX17 and FOXA2 (Tewary et al., 2019). Therefore, we used these two micropatterned differentiation protocols to evaluate the differentiation responses of all four hESC lines (**Fig. 2A**). After the 48-hour BMP4 induction, we observed different fate patterning between the lines. VUB01 and VUB03 showed the radially segregated peri-gastrulation-like fates, VUB02 colonies were mostly overtaken by T-positive cells, whereas VUB04 colonies only showed a slight upregulation of T at the very edge of the colonies **(Fig. 2B)**. Quantitative analysis confirmed that most of the cells within VUB04 colonies retained the expression of SOX2, whereas fewer than 10% were positive for T (**Fig. 2C**). Importantly, VUB04 also demonstrated a poor response in the DE micropatterned differentiation. Here again, we observed a similar variability in fate patterning as with the BMP4 induction, with VUB02 colonies largely consisting of cells positive for DE markers, whereas VUB04 colonies only displaying a thin outside ring of SOX17- and FOXA2-double positive cells (**Fig. 2D**). Although variability was observed in the amount of double positive cells between the colonies of the same line, VUB04 consistently showed the worst induction towards the DE fate (**Fig. 2E**). These results were consistent with our previous finding that VUB04 displays a low differentiation propensity towards DE in directed differentiation protocols when starting from monolayer cultures.

**Figure 2.**
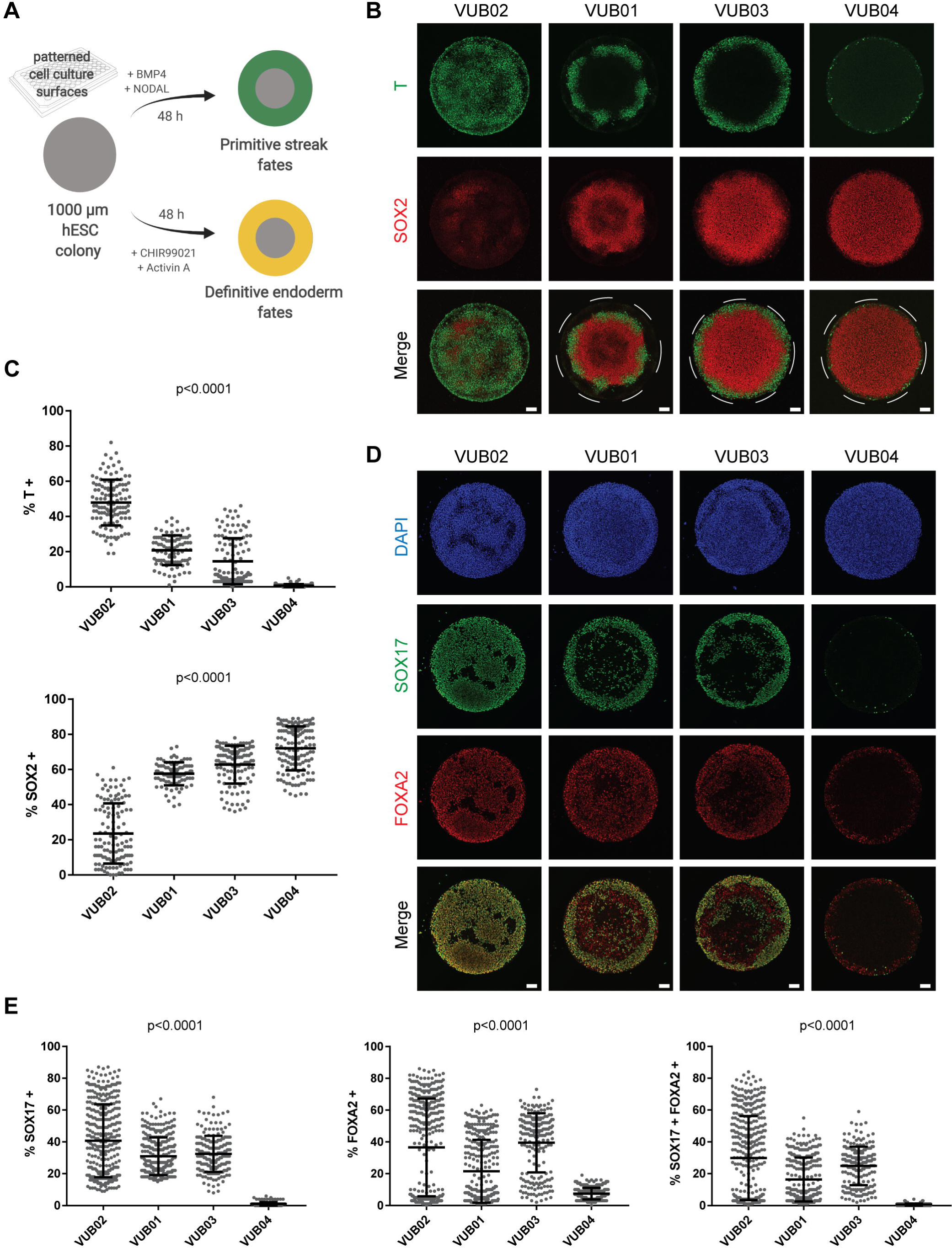
Micropatterned differentiation confirms the low differentiation propensity of VUB04 towards definitive endoderm (DE). A) Schematic overview of micropatterned differentiation protocols used to direct four hESC lines towards primitive streak and DE like fates in 1000 μm in diameter circular colonies. Created with BioRender. B) Representative immunofluorescent images for T and SOX2 after 48-hour micropatterned differentiation towards primitive streak fates. C) Percentage of T- and SOX2-positive cells after peri-gastrulation-like differentiation observed within the tested lines. Each data point represents one individual colony. Number of colonies were 120, 91, 120 and 134 for VUB02, VUB01, VUB03 and VUB04, respectively. Data pooled from two independent experiments. Error bars represent mean ± SD. D) Representative images for SOX17 and FOXA2 after 48-hour DE micropatterned differentiation. E) Percentage of SOX17-positive, FOXA2-positive and double-positive cells after DE differentiation observed in the tested lines. Each data point represents one individual colony. Number of colonies were 370, 318, 201 and 300 for VUB02, VUB01, VUB03 and VUB04, respectively. Data pooled from two independent experiments. Error bars represent mean ± SD. All p-values were calculated using one-way ANOVA. All scale bars represent 100 μm.

### Distinct transcriptomic profile of undifferentiated VUB04 cells does not explain the low DE differentiation propensity

Based on the observations described above, we hypothesised that VUB04 does not respond efficiently to DE differentiation cues because it differently regulates the activity of signalling pathways involved in DE specification. Therefore, we performed bulk mRNA-sequencing of the hESC lines at the undifferentiated stage to identify potential differences in the transcriptomic profile of VUB04. Unsupervised clustering analysis of the transcriptomic data showed that all VUB04 samples cluster separately from the other three lines (**Fig. 3A**). Although each hESC line demonstrated some differences in its expression profile, the first Principal Component which accounted for 41.7% variability between all the samples clustered all the VUB04 samples away from the other samples (**Fig. 3B**). Therefore, in the following differential expression analysis we compared VUB04 to the other three lines, grouped as the control. The analysis showed that 579 genes are more than twofold up- or down-regulated in VUB04 at the FDR < 0.05 significance level (**Fig. 3C, Fig. S2A**). Additionally, we observed differential expression of the pluripotency markers *POU5F1, SOX2* and *NANOG* in VUB04 (**Fig. 3D**). Although these were less than a twofold difference, it was an interesting observation given the high number of POU5F1-positive cells in VUB04 after 72h of DE differentiation **(Fig. 1E)**. For *NANOG*, which was the most differentially expressed, we performed a copy number assay to check for a possible *NANOG* gene duplication in VUB04. The result clearly showed that VUB04 has only two copies of the *NANOG* gene (**Fig. S2B**).

**Figure 3.**
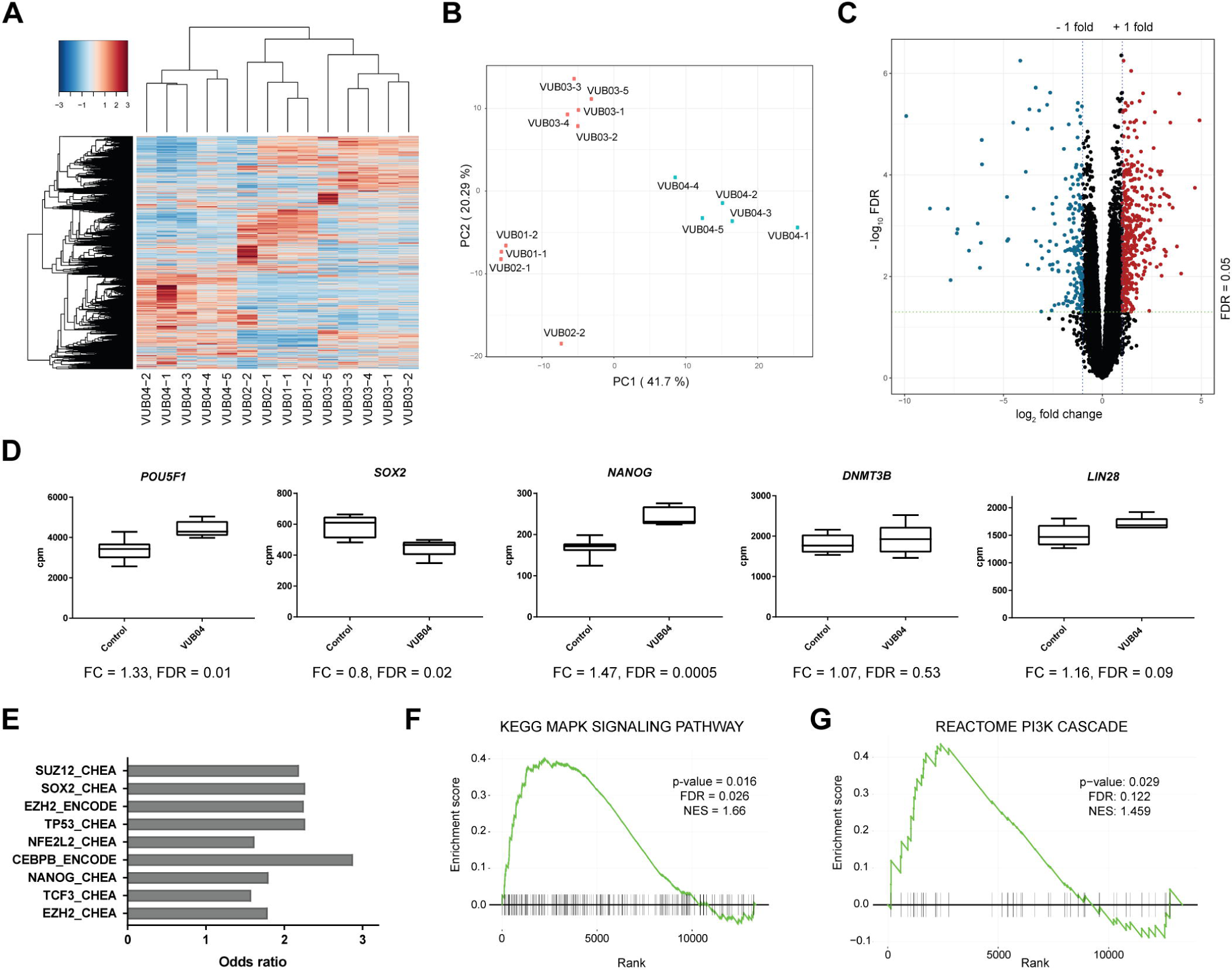
Transcriptomic analysis reveals that VUB04 has a distinct expression profile at the undifferentiated stage. A) Unsupervised hierarchical clustering and heatmap of transcriptome data of all lines tested. B) Principal Component Analysis of dimension 1 versus dimension 2 based on normalized transcriptome data. C) Volcano plot based on a comparison of gene expression levels between VUB04 and the control group (VUB01, VUB02 and VUB03). Genes with |log_2_ fold change| > 1 and FDR < 0.05 were marked as either significantly upregulated (red) or downregulated (blue). D) Normalized expression values for selected pluripotency markers in the control group and VUB04. FC = fold change. E) Transcription factor enrichment for the top deregulated genes in VUB04 (based on top |log_2_ fold change| and FDR < 0.05) done by the Enrichr tool and based on CHEA and ENCODE databases.) F) and G) Enrichment profile for MAPK/ERK signalling (F) and PI3K signalling (G) performed on the ranked gene list based on a comparison of the expression levels between VUB04 and the control group. Rank 0 represents the gene with the highest-ranking score. Vertical black bars represent genes within the ranked list belonging to the given pathway. NES = Normalized Enrichment score.

To evaluate which factors may contribute to the distinct VUB04 expression profile, we performed additional bioinformatic analysis. We carried out transcription factor enrichment analysis for the top differentially expressed genes in VUB04 using the Enrichr tool. Interestingly, the analysis showed that the list of top differentially expressed genes is significantly enriched for SOX2 and NANOG targets (**Fig. 3E**). We then checked if any signalling pathways associated with the pluripotent state are enriched within the significantly differentially expressed genes in VUB04. GSEA analysis identified MAPK/ERK signalling as one of the most enriched pathways within the most upregulated genes in VUB04 (**Fig. 3F, Fig. S2C**). The MAPK/ERK pathway together with PI3K/AKT signalling have been suggested to be induced by FGF2 in hPSCs and to play important roles in the maintenance of pluripotency (Lanner and Rossant, 2010). However, the enrichment for PI3K signalling within the most upregulated genes in VUB04 was not statistically significant (FDR > 0.05; **Fig. 3G**). Furthermore, the expression profile of VUB04 at the undifferentiated stage did not indicate any clear deregulation for Activin/NODAL and WNT pathways, which are crucially involved in DE specification. Together, the transcriptomic analysis at the undifferentiated stage did not provide a clear reason for the VUB04 differentiation impairment.

### Transcriptomic profile after 24-hour DE differentiation suggests inefficient activation of WNT signalling in VUB04

We thus hypothesised that the influence of the distinct expression profile of VUB04 on its DE differentiation propensity is mainly manifested once the line is exposed to specific differentiation signals. During the first 24h of DE differentiation, cells were incubated with both the WNT signalling activator CHIR and Activin A – a ligand of the Activin/NODAL pathway. As Activin/NODAL signalling is active in both pluripotency and during endoderm specification, whereas WNT signalling is stimulated only during the first 24h of endoderm differentiation and is required to switch the activity of Activin/NODAL pathway, we assessed whether WNT signalling was efficiently induced in VUB04 after this period. Immunofluorescent analysis showed that β-catenin is not present in the nuclei at the undifferentiated stage but changes its cellular localization after 24h DE differentiation in both VUB01 and VUB04 (**Fig. S3**). However, nuclear localization of β-catenin does not necessarily imply efficient expression of WNT signalling downstream targets. Therefore, we performed transcriptomic analysis at the 24-hour DE differentiation timepoint, using VUB01 and VUB02 as a control group, as they robustly differentiate towards mesendodermal derivatives (**Fig. 1, Fig. 2**), indicating efficient activation of WNT signalling. As in the undifferentiated state, VUB04 displayed a different expression profile, clustering away from the control lines (**Fig. 4A**). There were 1054 genes expressed twofold higher or lower in VUB04 samples than in the control lines at the FDR < 0.05 significance level (**Fig. 4B**). Transcription factor enrichment analysis indicated that the list of top deregulated genes in VUB04 is significantly enriched for NANOG and SOX2 (**Fig. 4C**), again suggesting an inefficient exit from pluripotency after 24-hour differentiation (**Fig. 4D**). While pluripotency genes remained upregulated, many downstream targets of the WNT pathway (e.g. *LEF1, DKK1, DKK4*) had little to no expression, and most of the genes associated with primitive streak formation (e.g. *MIXL1, EOMES, T, GSC*) were expressed at a much lower level than in the control group (**Fig. 4D**). In agreement, Ingenuity Pathway Analysis for upstream regulators predicted that the PI3K/AKT signalling, which is linked to maintenance of the pluripotent state, and GSK3, a negative regulator of the WNT pathway, are activated in the VUB04 samples in comparison to the control group (**Fig. 4E**).

**Figure 4.**
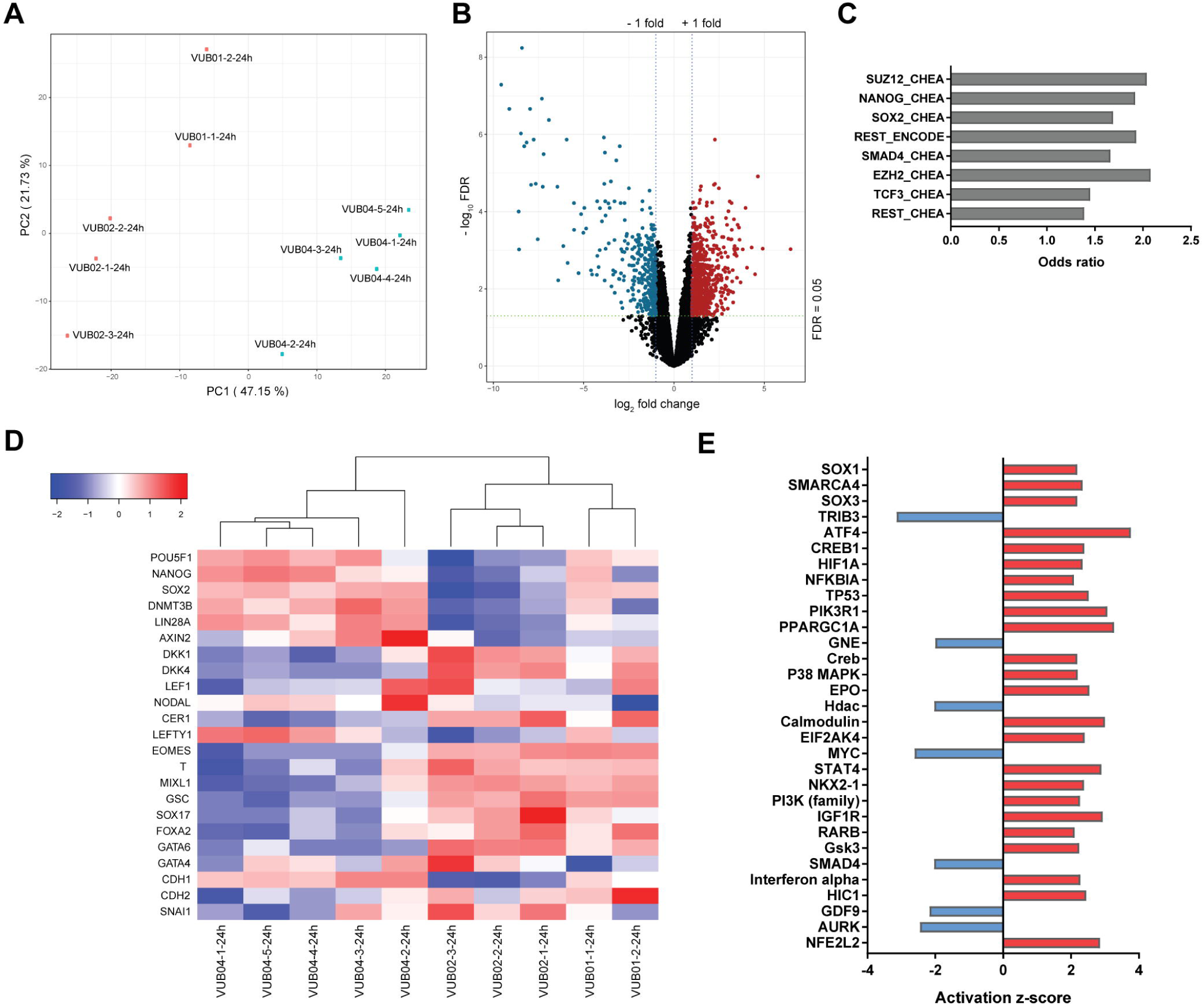
Transcriptomic profile after 24-hour DE differentiation suggests inefficient activation of WNT signalling in VUB04. A) Principal Component Analysis of dimension 1 versus dimension 2 based on normalized transcriptome data. B) Volcano plot based on comparison of gene expression levels between VUB04 and the control group (VUB01 and VUB02). Genes with |log2 fold change| > 1 and FDR < 0.05 were marked as either significantly upregulated (red) or downregulated (blue). C) Transcription factor enrichment for the top deregulated genes (|log_2_ fold change| and FDR < 0.05) in VUB04 done by Enrichr tool. D) Heatmap for gene expression levels (normalized counts per million) of pluripotency genes, WNT and NODAL signalling downstream targets, DE and epithelial-to-mesenchymal transition markers in the control group and in VUB04 after 24-hour DE differentiation. E) Prediction for upstream regulators of differentially expressed genes in the 24-hour DE VUB04 samples with |log_2_ fold change| > 1 and FDR< 0.05 done by Ingenuity Pathway Analysis. Only the upstream regulators with the activation z-score higher than 2 or lower than −2 and p-value < 0.05 are shown.

### Increased stimulation of WNT signalling improves VUB04 differentiation efficiency towards DE

As WNT signalling appeared to be inefficiently activated in VUB04 during the first 24h of DE differentiation, we modified the protocol in an attempt to rescue the poor differentiation of VUB04. We observed an increase in SOX17-positive cells in VUB04 when stimulated with WNT inducer CHIR at concentrations higher than the standard 3 µM (**Fig. 5A, Fig. S4A**), with an almost 5-times higher expression of *SOX17* and *FOXA2* when treated with 9 µM CHIR (**Fig. 5B**). The differentiation outcome also improved when the 3 µM CHIR condition was supplemented with the PI3K inhibitor LY294002 (LY) for the first 48h of differentiation, (**Fig. 5 A, B, Fig. S4A**). Intriguingly, the expression levels of pluripotency markers in VUB04 only decreased to similar levels as in VUB01 when the cells were treated with LY, but not when higher concentrations of CHIR were used (**Fig. 5B, Fig. S4B**). Additionally, higher CHIR concentrations seemed to increase the expression of the mesodermal markers *PDGFRA* in both lines tested and *KDR* in VUB04 (**Fig. S4B**). These results suggest that active PI3K/AKT signalling may prevent efficient activation of WNT signalling at the onset of DE differentiation (**Fig. 5C**), as previously suggested (Singh et al., 2012).

**Figure 5.**
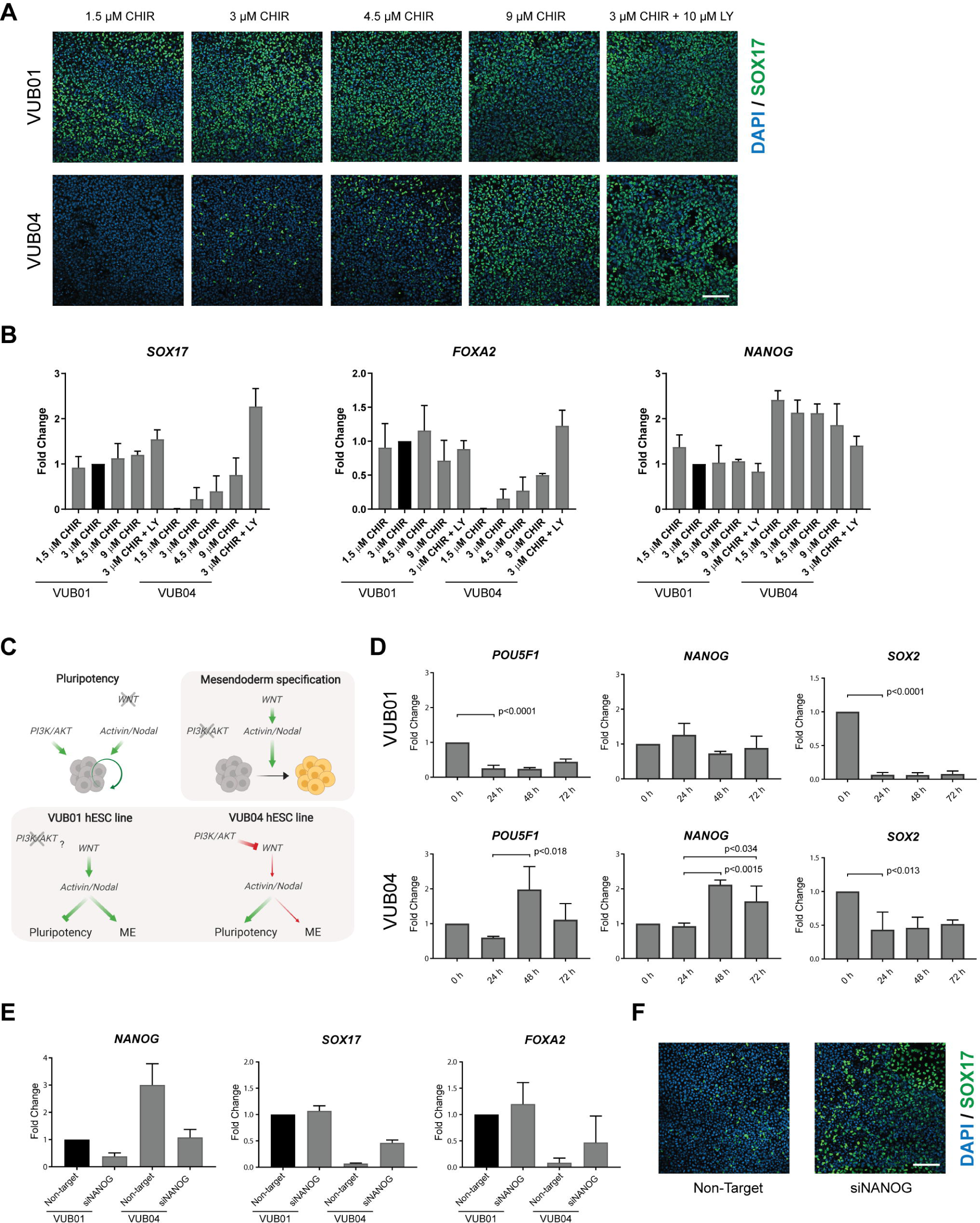
Stronger activation of WNT signalling improves DE differentiation efficiency of VUB04. A) Representative immunofluorescent images for SOX17 after 72-hour DE differentiation of VUB01 and VUB04 in various differentiation conditions. The scale bar represents 100 μm. B) Gene expression analysis of *SOX17, FOXA2* and *NANOG* in VUB01 and VUB04 differentiated for 72h towards DE – standard differentiation condition (3µM CHIR99021 for the first 24h) was compared to modified conditions (different concentration of CHIR99021 during the first 24h or additional incubation with PI3K inhibitor LY294002 for the first 48h). Data represents three biological replicates. C) Schematic illustration of the signalling pathways involved in the maintenance of pluripotency (white background) and mesendoderm (ME) specification (grey background). The lower part illustrates the difference in signalling pathway crosstalks between VUB01 and VUB04 hESC lines which influence the differentiation outcome. Created with BioRender. D) Gene expression dynamics of pluripotency markers in VUB01 and VUB04 during the 72-hour DE differentiation. Data represents three biological replicates. The p-values were calculated with unpaired t-test. E) Comparison of *NANOG, SOX17* and *FOXA2* expression after 72-hour DE differentiation between VUB01 and VUB04 transfected cells. Data represents two biological replicates. F) Representative images for SOX17 after 72-hour DE differentiation of VUB04 transfected with either non-targeting or NANOG-targeting siRNA. The scale bar represents 100 μm.

### *NANOG* expression dynamics during differentiation define DE specification efficiency

As the data indicated that VUB04 did not efficiently exit the pluripotent state during the DE differentiation, we compared the dynamics of pluripotency factor expression between VUB01 and VUB04 over the course of the 72-hour unmodified DE differentiation protocol. In VUB01, the expression of *POU5F1* and *SOX2* significantly decreased after 24h, whereas *NANOG* expression levels remained similar to the undifferentiated stage during the entire 72h differentiation period (**Fig. 5D**). VUB04 displayed a different pattern: after 24h the downregulation of *SOX2* was less pronounced and *POU5F1* and *NANOG* expression increased significantly between 24h and 48h. The latter observation coincides with CHIR withdrawal and subsequent incubation with Activin A only for the following 2 days (**Fig. 5D**). Under normal circumstances, 24h WNT induction should result in the shift of Activin/NODAL signalling activity from promoting pluripotency to driving DE specification. The fact that *POU5F1* and *NANOG* expression levels increased at the 48-hour timepoint indicated that exit from pluripotency is not efficiently induced in VUB04, likely due to the impaired activation of WNT signalling, and that the cells remain programmed to support pluripotency when exposed to a high concentration of Activin A (**Fig. 5C**).

NANOG is one of the key nodes of the pluripotency network (Boyer et al., 2005) and it cooperates together with Activin/NODAL signalling in supporting hPSC self-renewal (Brown et al., 2011). As inhibition of PI3K/AKT signalling in VUB04 seemed to improve the exit from pluripotency and DE differentiation, we explored if reducing *NANOG* expression in VUB04 prior to DE differentiation would also improve the differentiation outcome. Interestingly, knocking down *NANOG* led to improved DE differentiation (**Fig. 5E, F, Fig. S5B**), with increased expression of *SOX17* and *FOXA2* after 72h, though the expression levels were not as high as in VUB01 (**Fig. 5E**). Nevertheless, *NANOG* knock down in VUB04 also led to improved HP differentiation (**Fig. S5C**). At the same time, exogenous downregulation of *NANOG* in VUB01 did not result in any change in its high DE differentiation capacity (**Fig. 5E, Fig. S5B**). Therefore, these experiments demonstrated that exogenous knockdown of *NANOG* could eliminate the failure of VUB04 to differentiate to DE. It also suggested that efficient DE specification is dependent on a certain level of *NANOG* expression, with lower expression levels likely circumventing the tempering effect that active PI3K/AKT pathway has on WNT signalling at the onset of differentiation (**Fig. 5C**).

## Discussion

Functional variability among hPSC lines is a significant obstacle for their efficient use in many biomedical applications (Keller et al., 2018; Ortmann and Vallier, 2017). Here, we used two standardized short-term differentiation assays, conventional monolayer and micropatterned differentiation, to screen hESC lines for their differentiation efficiency towards DE and subsequently evaluated how differences in the intrinsic regulation of signalling pathways influence the differentiation outcome. Importantly, we obtained the same predictions for DE differentiation efficiencies when the same hESC lines were subjected to these two differentiation assays in two different laboratories. We thus confirmed our recent report (Tewary et al., 2019) by showing that peri-gastrulation-like fate patterning can be used efficiently to fingerprint hPSC lines for their differentiation potential in a standardized, quantitative and robust manner. In agreement with previous studies (Hu et al., 2010; Kattman et al., 2011), we also demonstrated that it is possible to improve the low differentiation propensity of an individual line by optimising the differentiation conditions.

In the differentiation propensity screen, we identified VUB04, as a hESC line with a very low differentiation efficiency towards DE. Gene expression profiling during DE differentiation showed that VUB04 had a distinct expression dynamics of pluripotency genes which was coupled with inefficient priming for differentiation upon stimulation of Activin/NODAL and WNT signalling. Thus, VUB04 is a case in point that some line-specific properties can cause low differentiation efficiency towards DE fates already at the very onset of differentiation. The necessity to redirect the Activin/NODAL pathway activity to DE fate specification by WNT signalling (D’Amour et al., 2006; Funa et al., 2015; Yoney et al., 2018) is a vulnerable spot as any disruption of this process can result in a line with low DE efficiency.

Our study also suggests that active PI3K/AKT signalling has a tempering effect on efficient activation of WNT signalling during DE differentiation, as inhibiting PI3K/AKT signalling led to increased DE efficiency in VUB04. The mechanism by which PI3K/AKT signalling regulates pluripotency and early lineage specification is still being investigated. One study proposed that it plays a central role in maintaining hPSC pluripotency by modulating Activin/NODAL signalling to support self-renewal and by suppressing MAPK/ERK and WNT signalling pathways to prevent mesendoderm specification (Singh et al., 2012). According to this model, the switch in the Activin/NODAL signalling activity requires inactivation of the PI3K/AKT pathway, which in turn allows the MAPK/ERK signalling to inhibit GSK3β and subsequently to stimulate the WNT signalling to initiate differentiation. Although both the MAPK/ERK pathway and PI3K/AKT signalling were previously reported to be downstream targets of FGF2 signalling in hPSCs, the authors propose that in culture conditions supportive of pluripotency, PI3K/AKT signalling maintains the activity level of the MAPK/ERK signalling within the range that supports the undifferentiated state (Singh et al., 2012). A very recent study reported that PI3K/AKT pathway is active in early human embryos and showed that hPSCs could be expanded *in vitro* in the presence of PI3K activators and Activin A without the addition of FGF2 (Wamaitha et al., 2020), which may suggest a dominant role of PI3K pathway over MAPK/ERK signalling in supporting hPSC self-renewal. Based on these studies and our data, we suspect that both the activity of MAPK/ERK and PI3K/AKT are elevated in VUB04 with PI3K signalling playing a dominant role in preventing VUB04 from exiting the pluripotent state during DE differentiation. Additional proteomic analysis of the PI3K/AKT and MAPK/ERK components would provide more evidence for this signalling pathway crosstalk.

Interestingly, reducing *NANOG* expression at the undifferentiated stage in VUB04 also improved DE specification. NANOG has been shown to be necessary during mesendoderm specification, whereas its strong downregulation initiates neuroectoderm specification (Vallier et al., 2009; Wang et al., 2012). Thus, our results suggest that the effect of NANOG on hPSC differentiation depends on its expression level, as downregulation of *NANOG* in the control line, VUB01, was still permissive for mesendoderm differentiation and did not result in any reduction in DE differentiation efficiency. Additionally, it is likely that the exogenous downregulation of *NANOG* in VUB04 directly or indirectly influenced the activity of signalling pathways, one of which is possibly PI3K/AKT, which otherwise suppressed efficient activation of WNT signalling.

To conclude, this study presents relevant insight into the influence of WNT and PI3K/AKT signalling on hPSC differentiation towards DE and demonstrates that a hPSC line with a specific differentiation impairment can be used as a tool to study crosstalks between signalling pathways involved in the early lineage specification.

## Experimental procedures

### Human ESC culture

Human ESC lines VUB01, VUB02, VUB03 (VUB03_DM1) and VUB04 (VUB04_CF) were derived and characterized as previously described (Mateizel et al., 2006). Cells were routinely cultured on dishes coated with 5 µg/ml LN521 (Biolamina) in NutriStem® hESC XF medium (NS medium; Biological Industries) with 100 U/ml Penicillin-Streptomycin (Pen/Strep; Thermo Fisher Scientific) and passaged as single cells in a 1:10 to 1:30 ratio using TrypLE™ Express (Thermo Fisher Scientific) when 70-80% confluent. The cells were kept at 37°C in 5% CO2. All hESC lines were analysed for their genetic content by array-based comparative genomic hybridization (Human Genome CGH Microarray 4×44K, Agilent Technologies) as previously described (Jacobs et al., 2014), and chromosomally balanced frozen bulks were prepared prior to the onset of experiments. Karyotypes and passage numbers of lines used in the study can be found in **Table S1**.

### Directed differentiation protocols

#### Mesendoderm and definitive endoderm specification

We used a modified protocol based on D’Amour *et al*. and Sui *et al*. adapted to LN521 coating (D’Amour et al., 2005; Sui et al., 2012). Briefly, hESCs were seeded at a density of 4 × 10^4^ cells per cm^2^ and the differentiation was started 1-2 days later once the cells were 50-60% confluent. Cells were differentiated for 24h in RPMI 1640 Medium with GlutaMAX™ and supplemented with 2% B27 supplement (both from Thermo Fisher Scientific), 3 µM CHIR99021 (CHIR, STEMCELL Technologies) and 100 ng/mL Activin A (Biotechne). After the 24-hour mesendoderm specification, the cells were incubated for an additional 48h in the same differentiation medium but without CHIR. When indicated, the DE differentiation conditions were modified: either 1.5, 3, 4.5 or 9 µM CHIR was used during the first 24h, or the CHIR concentration was left at 3 µM but 10 µM of PI3K inhibitor LY294002 was added to the differentiation medium for the first 48h.

#### Hepatic progenitors

The protocol was adapted from Cameron *et al*. (Cameron et al., 2015). Human ESCs were seeded on LN521 at a density of 4 × 10^4^ cells per cm^2^. The next day, the pluripotency medium was changed to RPMI 1640 medium supplemented with GlutaMAX™, 0.5% B27 supplement, 3 µM CHIR and 100 ng/mL Activin A. After the first 24h of differentiation CHIR was removed. The day after, the medium was changed to KnockOut™ DMEM containing 20% KnockOut™ Serum Replacement, 0.5% GlutaMAX™ supplement, 1% MEM Non-Essential Amino Acids, 100 U/ml Pen/Strep (all from Thermo Fisher Scientific), 0.1 mM β-mercaptoethanol and 1% DMSO (both from Sigma-Aldrich). This differentiation step lasted until day 8 and the medium was refreshed daily.

Details on differentiation towards neuroectoderm lineage and EB differentiation can be found in the Supplemental Information.

### Micropatterned differentiation protocols

Microtiter 96-well plates with patterned islands of 1000 µm in diameter were prepared following a previously published protocol (Tewary et al., 2017). Prior to seeding cells onto the plates, the wells were activated with N-(3-Dimethylaminopropyl)-N′-ethylcarbodiimide hydrochloride and N-Hydroxysuccinimide (both from Sigma-Aldrich) for 20 minutes. The plates were thoroughly triple washed with ddH_2_O and incubated with 10 µg/mL LN521 diluted in Dulbecco’s phosphate-buffered saline (DPBS; Thermo Fisher Scientific) with calcium and magnesium for 3 hours at 37°C or at 4°C overnight. After incubation, the plates were triple washed with DPBS to remove any passively adsorbed extracellular matrix.

To develop micropatterned colonies, hESCs were seeded in NS medium with 10 µM ROCK inhibitor (ROCKi) Y-27632 at a density of 8 x 10^4^ cells per well and incubated for 2 h at 37°C. Although ROCKi is not essential for seeding hESCs as single cells on LN521, its addition to the seeding suspension yielded better long-term attachment of micropatterned hESC colonies. After 2-3 h, the medium was changed to NS without ROCKi. When confluent colonies were observed, typically 12-18 h after seeding, induction towards primitive streak associated and DE fates was started. For both differentiation protocols the basal medium was N2B27 consisting of 93% DMEM, 1% Pen/Strep, 1% MEM Non-Essential Amino Acids, 0.1 mM β-mercaptoethanol, 1% GlutaMAX™, 1% N2 Supplement and 2% B27 minus retinoic acid supplement (all from Thermo Fisher Scientific). To generate gastrulation associated fates, the N2B27 medium was supplemented with 50 ng/mL BMP4, 100 ng/mL NODAL and 10 ng/mL FGF2 (all from Biotechne). For the induction of the definitive endoderm associated fates, N2B27 medium was supplemented with 3 µM CHIR (STEMCELL Technologies) and 100 ng/mL Activin A (Biotechne). After 48h of differentiation, the micropatterned colonies were fixed for subsequent protein expression analysis.

### Immunofluorescent stainings and image analysis

For immunofluorescent stainings, the cells were fixed with 3.7% paraformaldehyde (Sigma-Aldrich) for 20 min, rinsed three times with DPBS and then permeabilized with 100% methanol (Sigma-Aldrich) for 3 min. Blocking was performed using 10% Fetal Bovine Serum (FBS; Thermo Fisher Scientific) in DPBS at 4°C overnight. Primary antibodies were diluted in 10% FBS and incubated at 4°C overnight. Then, cells were triple rinsed with DPBS and incubated with secondary antibodies and 10 µg/mL Hoechst 33342 diluted in DPBS with 10% FBS at room temperature for 2h, followed by the final triple rinse with DPBS. The nuclei in micropatterned hESC colonies were stained with DAPI instead of Hoechst. Antibody sources and concentrations are shown in **Table S2**.

Immunofluorescent images were taken using a LSM800 confocal microscope (ZEISS). The cell count analysis presented in Fig. S4 was done using ZEN desk imaging software (ZEISS). The immunofluorescent images for quantitative DE differentiation data presented in Fig. 1E were taken using an IX-81 fluorescent microscope (Olympus) with Cell^F software (Olympus) and counted with ImageJ software. We analysed at least 1000 cells per biological replicate. To obtain quantitative single-cell data from micropatterned colonies the plates were scanned with Cellomics Arrayscan VTI platform (Thermo Fisher Scientific) using the ‘TargetActivation.V4’ bioassay algorithm. This algorithm utilizes the fluorescent intensity in the DAPI channel to identify individual nuclei in all fields imaged and acquires the associated intensity of proteins of interest localized within the identified region. Single-cell data extracted from immunofluorescent images were exported into a custom-built software for image analysis, ContextExplorer (Ostblom et al., 2019), which classifies cells into colonies and calculates the percentage of cells positive for proteins of interest per single colony.

### Quantitative real-time PCR analysis

For qRT-PCR gene expression analysis, total RNA was extracted using the RNeasy Mini Kit or RNeasy Micro Kit (Qiagen) with on-column DNase digest. Reverse transcription was performed using the First-Strand cDNA Synthesis Kit (GE Healthcare). Quantitative RT–PCR was performed using qPCR MasterMix Plus Low ROX (Eurogentec) and TaqMan Gene Expression Assays (Thermo Fisher Scientific). The samples were run on the ViiA 7 thermocycler (Thermo Fisher Scientific) using standard cycling parameters provided by the manufacturer. The relative expression of genes of interest was calculated by the ΔCt and ΔΔCt method with *GUSB* used as a reference gene. For gene expression analysis after the differentiation towards hepatic progenitors, *UBC* was used as a second reference gene. References for the TaqMan assays can be found in **Table S3**.

### Statistical analysis

Statistical analysis for quantitative immunofluorescent data and qRT-PCR gene expression analysis was performed using the GraphPad Prism software (version 7.04; GraphPad Software, Inc.). An unpaired t-test was performed when comparing two conditions and a one-way ANOVA followed by Bonferroni correction for more than two conditions. The significance level was set at p-value <0.05.

### mRNA sequencing

Total RNA was extracted using the RNeasy Mini Kit (Qiagen) with on-column DNase digest. The concentration and quality of extracted RNA were evaluated using the Quant-iT RiboGreen RNA Assay Kit (Thermo Fisher Scientific) and the RNA 6000 Pico Chip (Agilent Technologies), respectively. Subsequently, 500 ng of RNA was used to perform an Illumina sequencing library preparation using the QuantSeq 3’ mRNA-Seq Library Prep Kit (Lexogen) according to the manufacturer’s protocol. During library preparation 15 PCR cycles were used. Libraries were quantified by qRT-PCR, according to Illumina’s protocol ‘Sequencing Library qPCR Quantification protocol guide’, version February 2011. The library’s size distribution and quality were analysed using the High sensitivity DNA Chip (Agilent Technologies) on a 2100 Bioanalyzer platform (Agilent Technologies). Sequencing was performed on a NextSeq 500 (Illumina), generating 75 bp single-end reads. Two separate sequencing runs where performed, one for the undifferentiated hESC samples and one for the 24-hour ME differentiation samples. All data were deposited in the GEO repository with accession number GSE148050.

Details on the downstream bioinformatic analysis of mRNA sequencing can be found in the Supplemental Information.

### Transfection for siRNA knockdown

Human ESCs were seeded on LN521 at a density of 4 × 10^4^ cells per cm^2^ and the transfection was done the next day once the cells were around 50% confluent. The siRNAs used were ON-TARGETplus Human NANOG siRNA SMARTpool and ON-TARGET*plus* Non-targeting siRNA (Dharmacon, Cat.No L-014489-00-0005 and D-001810-01-05, respectively). Transfection was performed using siRNAs at the final concentration of 50nM and Lipofectamine RNAiMAX diluted in OptiMEM medium following manufacturer’s protocol (Thermo Fischer Scientific). The cells were incubated with transfection reagents for 24h after which the differentiation was started.

## Author Contributions

D.D. co-designed the project, performed most experiments and co-wrote the manuscript, M.T. helped with micropatterned differentiation experiments and protein expression analysis, A.K. helped with the hepatic differentiation experiments, L.T. and F.V.N. performed the RNA sequencing and most bio-informatic analyses, L.P. assisted with transfection experiments, J.O. helped with micropatterned differentiation analysis, E.C.D.D. performed part of bio-informatic analyses, C.M. and S.F. helped with protein expression analysis, C.S. and P.W.Z. provided important intellectual contributions, K.S. supervised the project and co-wrote the manuscript, M.G. supervised the project, co-designed and co-wrote the manuscript. All authors revised and approved the manuscript.

## Acknowledgments

This work was supported by the Methusalem grant of Vrije Universiteit Brussel granted to K.S. D.D. is a PhD fellow of Research Foundation – Flanders (Fonds voor Wetenschappelijk Onderzoek, FWO – Vlaanderen). A.K. is a PhD fellow of FWO (Strategisch Basisonderzoek). The authors would like to thank Geoffrey Duque for the technical assistance with LSM800 confocal microscope and the NXTGNT team for performing the mRNA-sequencing.

