## Supplemental information for "Endogenous suppression of WNT signalling in human embryonic stem cells leads to low differentiation propensity towards definitive endoderm"

#### Supplemental Figures

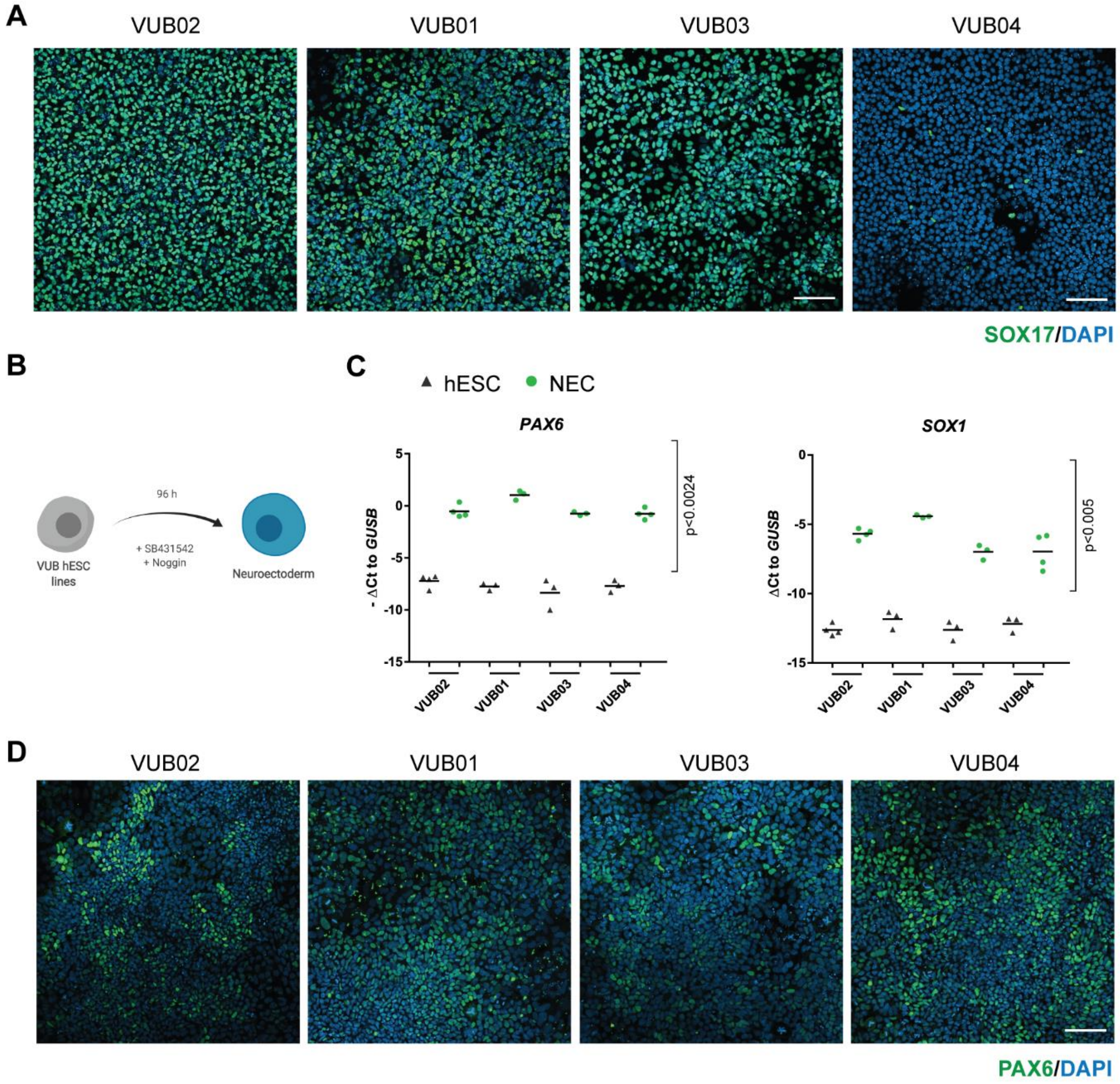


**E**

| Sample Name | Self-renewal | Ectoderm | Mesoderm | Endoderm |
|---|---|---|---|---|
| EB VUB02 | -3.60 | 0.73 | 3.65 | 0.64 |
| EB VUB01 | -4.30 | 1.69 | 3.03 | 0.74 |
| EB VUB03 | -2.32 | 1.58 | 0.42 | -0.42 |
| EB VUB04 | -4.37 | 0.19 | 5.74 | 0.61 |

Gene expression relative to the reference standard

| Upregulated | | | | | | Downregulated |
|---|---|---|---|---|---|---|
| x > 1.5 | 1.0 < x ≤ 1.5 | 0.5 < x ≤ 1.0 | -0.5 ≤ x ≤ 0.5 | -1.0 ≤ x < -0.5 | -1.5 ≤ x < -1.0 | x < -1.5 |

**Figure S1.** Differentiation propensity screen of all four hESC lines showing that VUB04 has a low differentiation efficiency towards DE but differentiates with the same efficiency towards neuroectodermal lineage. (A) Representative immunostainings for SOX17 after DE differentiation of all lines tested. (B) Schematic overview of the neuroectoderm (NEC) differentiation protocol used in the study. Created with BioRender. (C) Gene expression level of *PAX6* and *SOX1* in hESCs and NEC samples. Data was collected from at least three biological replicates and p-values were calculated using one-way ANOVA. (D) Representative immunofluorescent images for PAX6 after 96-hour NEC differentiation. All scale bars represent 100 µm. (E) The Scorecard assay result for equally-sized 12-day embryoid bodies spontaneously differentiated in serum-free APEL™ medium - gene expression levels of pluripotency markers and trilineage differentiation associated genes were compared to the Scorecard reference profile.

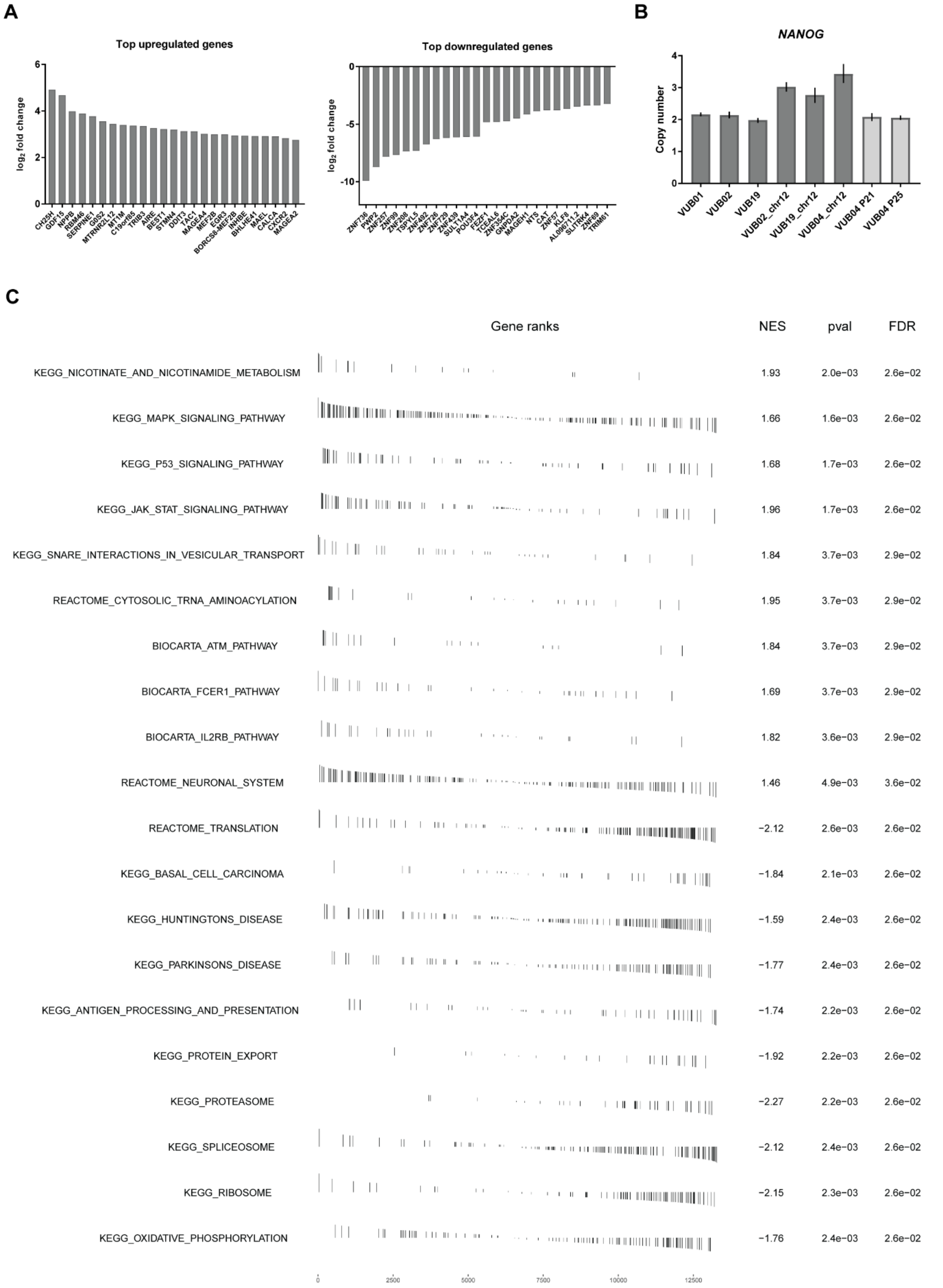


**Figure S2.** Global gene expression comparison between the control group (VUB01, VUB02 and VUB03) and VUB04 at the undifferentiated stage. A) Top 25 up- and down-regulated genes in VUB04 when

|$log_2$ fold change| > 1 and FDR < 0.05. B) Copy number assay for *NANOG*. Karyotypically normal sample of VUB19 hESC line was used as copy number = 2 control. Samples of VUB02_chr12, VUB19_chr12 and VUB04_chr12 hESC lines were known sublines carrying duplications on chromosome 12 which contain the *NANOG* gene locus. VUB04 P21 and VUB04 P25 are samples from VUB04 passages which were used during this study. The bar chart represents mean with range. C) Top 20 the most significantly enriched (FDR < 0.05) pathways in GSEA done on ranked gene list. The analysis was done using KEGG, Reactome and Biocarta pathway libraries. NES = Normalized Enrichment score.

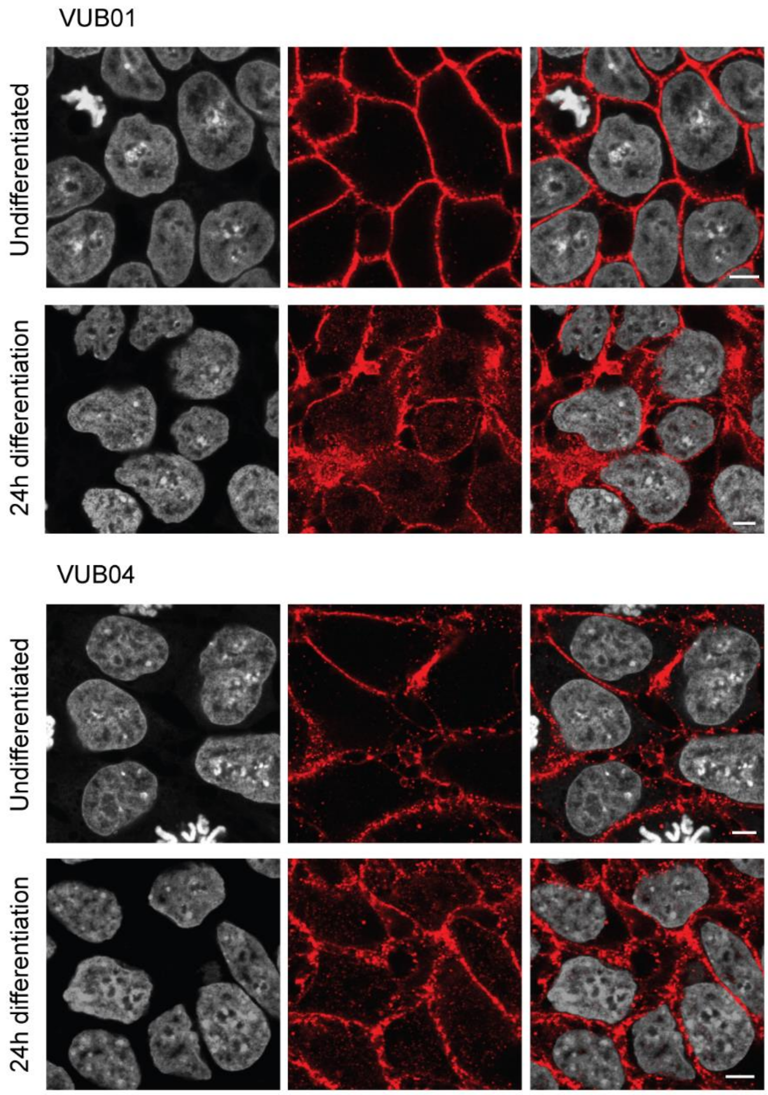


**Figure S3.** WNT signalling activation after the 24-hour DE differentiation. Representative immunofluorescent images for β-catenin in undifferentiated and differentiated VUB01 and VUB04. The scale bar represents 5 µm.

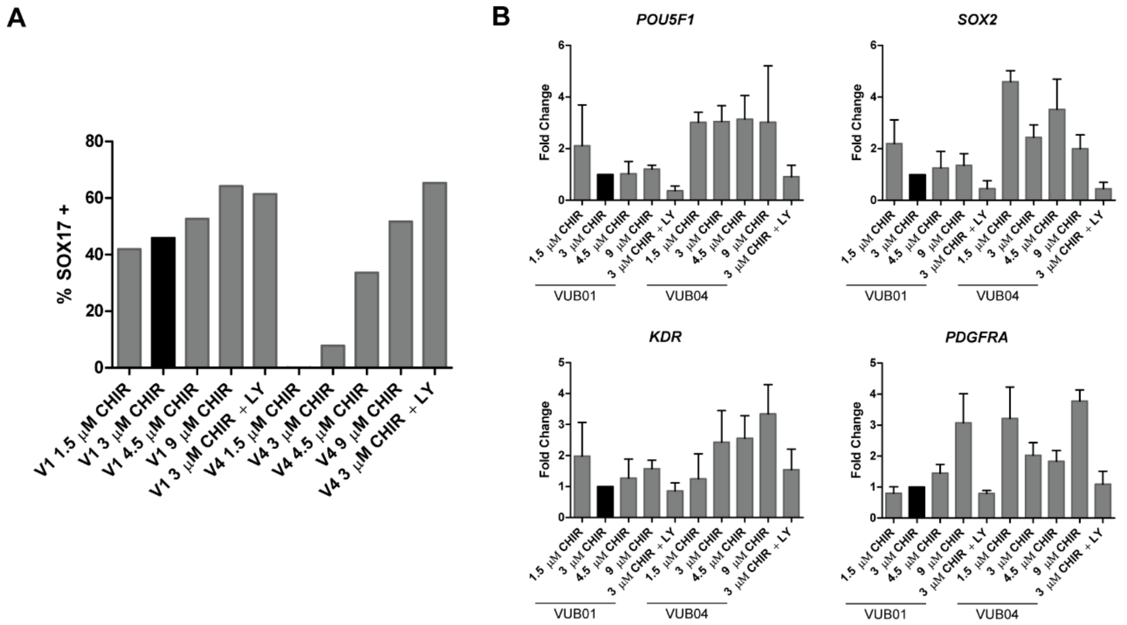


**Figure S4.** A) Percentage of SOX17-positive cells after the 72-hour DE differentiation of VUB01 and VUB04 in various differentiation conditions. Data represents one biological replicate. B) Expression of the pluripotency (*POU5F1*, *SOX2*) and mesoderm (*KDR*, *PDGFRA*) markers in DE samples of VUB01 and VUB04 after stronger activation of WNT signalling. The standard differentiation condition (3µM CHIR99021 for the first 24 h) was compared to modified conditions (different concentration of CHIR99021 during the first 24 h or additional incubation with PI3K inhibitor LY294002 for the first 48 h). Data represents three biological replicates.

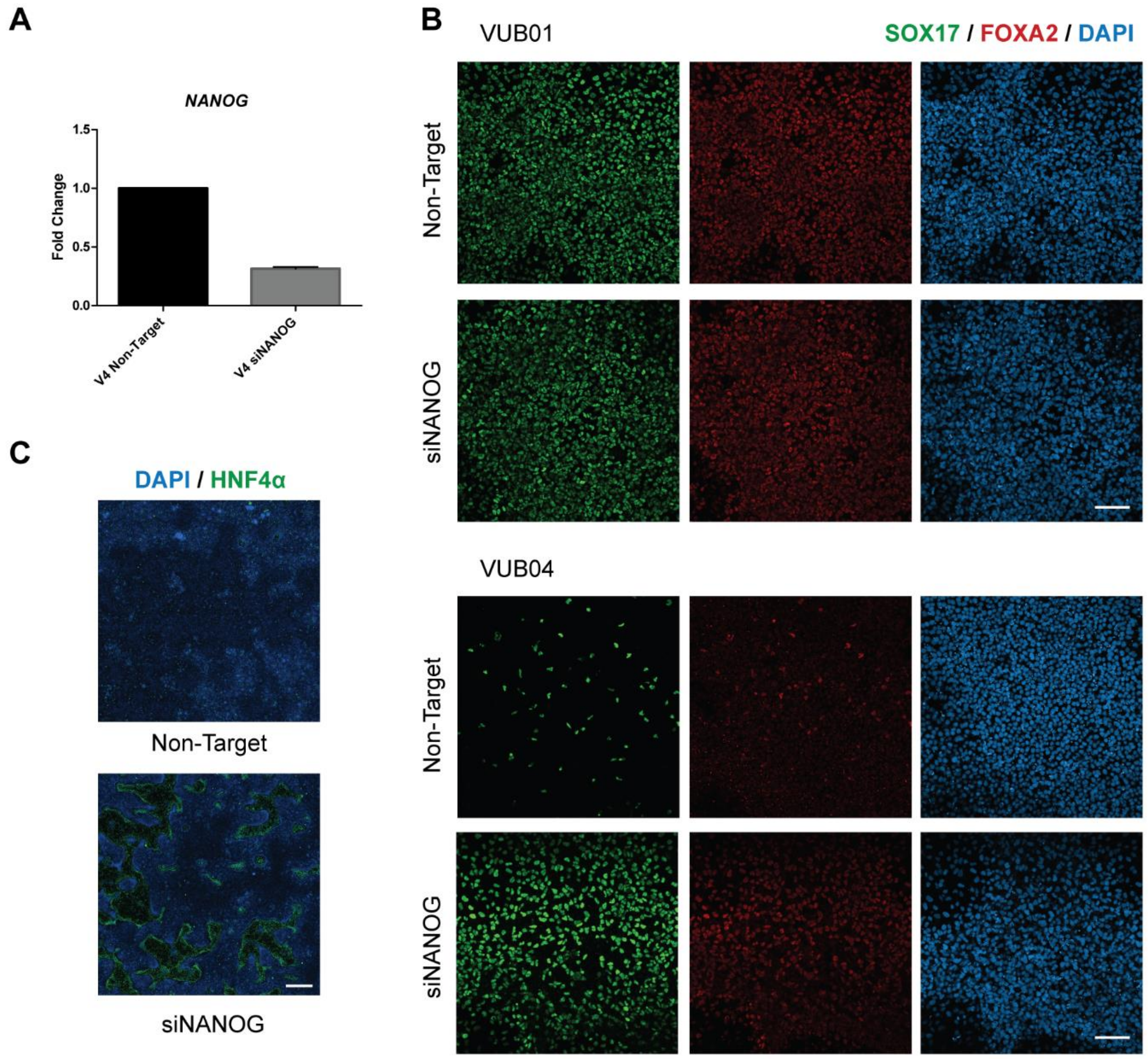


**Figure S5.** A) Comparison of *NANOG* expression level between VUB04 cells transfected for 24h with non-targeting siRNA and those transfected with siRNA against *NANOG*. Data represents three biological replicates. B) Knockdown of NANOG expression in VUB01 and VUB04 at the onset of DE differentiation. Representative immunofluorescent images for SOX17 and FOXA2 after 72-hour DE differentiation of VUB01 and VUB04 transfected with either non-targeting or *NANOG*-targeting siRNA. The scale bar represents 100 µm. C) Immunofluorescent images of whole well scans of VUB04 cells stained for HNF4α after transfection with *NANOG*-targeting siRNA followed by hepatic differentiation. The scale bar represents 1mm.

### Supplemental experimental procedures

#### Neuroectoderm specification

A widely used neural differentiation protocol was adapted to LN521-based culture (Chambers et al., 2009). Briefly, hESCs were seeded on LN521 at a density of 6 x $10^4$ cells per $cm^2$ and the differentiation was started 1-2 days later once the cells were 80-90% confluent. Cells were differentiated for 96 h in KnockOut™ DMEM medium containing 10% KnockOut™ Serum Replacement, 500 ng/mL Noggin (Biotechne) and 10 µM SB431542 (Biotechne). The medium was changed daily.

#### Embryoid body differentiation and gene expression analysis

Cells were passaged 2 days prior to the EB formation and harvested at 60-80% confluence. Single cell suspensions were generated using TrypLE™ Express and resuspended in STEMdiff™ APEL™ medium (STEMCELL Technologies) with 10 µM ROCKi. Cells were seeded at 5000 cells/well in the round-bottom non-treated 96-well plates (Corning® #3788) and spun down at 600 g for 5 min. EB differentiation lasted for 12 days and the medium was changed every third day. Total RNA extraction and cDNA conversion was performed as described for the qRT-PCR analysis in the main Methods section. The gene expression analysis was performed using the 384-well TaqMan™ hPSC Scorecard™ Panel (Thermo Fisher Scientific), which was run on the Viia7 thermocycler following the manufacturer's protocol.

#### mRNA sequencing

##### *Sequencing data analysis*

Sequencing quality was checked based on the parameters given on BaseSpace (Illumina). The quality of the raw reads was checked using the FastQC software tool, version 0.11.7, and the FastQ Screen software tool, version 0.11.4, was used to detect potential contamination. All samples had a similar sequencing depth with 19.7 x $10^6$ ± 3.4 x $10^6$ reads per sample. Trimming of library preparation adaptor sequences and poor-quality bases was performed by cutadapt (Martin, 2011) version 1.15. The high-quality reads were mapped to the Homo sapiens GRCh38.89 reference genome using STAR read mapper (Dobin et al., 2013), version 2.5.3a. To quantify the reads per gene, the RSEM software (Li and Dewey, 2011), version 1.3.0, was used. The estimated relative abundances were rounded off and grouped in a count table, one for each mRNA-sequencing run. Only coding genes were considered for further analysis.

The count tables were normalised using edgeR standard normalisation method (Robinson et al., 2010). To detect outlier samples in each mRNA sequencing experiment before differential expression analysis, clustering and principal component analysis (PCA) plots were built. This analysis only used genes which had a counts-per-million (cpm) above 1 in at least 2 samples. The PCA was performed on rlog-transformed counts (Love et al., 2016) using the R statistical computing software. A heatmap with the genes of the samples was made with normalised counts rescaled between -3 and 3 and the clustering was based on the Pearson correlation.

Differential gene expression analysis was performed using edgeR (Robinson et al., 2010) and only genes with a cpm greater than 1 in at least 5 samples within each mRNA-sequencing run were considered. For each comparison, a separated normalization was done with edgeR standard normalisation method and a different general linear model was built. Statistical testing was done using the empirical Bayes quasi-likelihood F-test. For multiple testing correction, the False Discovery Rate (FDR) method with the Benjamini-Hochberg procedure was used. Genes having FDR < 0.05 and a fold change > 2 or a fold change < 0.5 were considered as significantly differentially expressed in all downstream bioinformatic analyses.

*Enrichr analysis*

Transcription factor enrichment analysis was done using Enrichr (Chen et al., 2013; Kuleshov et al., 2016). The deregulated genes were used as input (|$\log_2$ fold change|>1, FDR<0.05). The outputs were selected from ENCODE and ChEA Consensus TFs from ChIP-X library with FDR < 0.05.

*Ingenuity pathway analysis*

Ingenuity Pathway Analysis (IPA; QIAGEN) was used for the upstream regulators analysis based on the differential gene expression between groups (Krämer et al., 2014). The data were uploaded with their respective $\log_2$ fold change, FDR and p-value. IPA predicts the activation state of regulators by correlating effects reported in literature with observed gene expression. In order to predict if a regulator is activated or inhibited, it computes the z-score. A z-score > 2 means activated, z-score < -2 means inhibited. Each z-score is associated with a p-value. The regulators classified as “chemical”, “complex”, “enzyme” and “other” were not included in the final plots.

*Gene set enrichment analysis*

The Gene set enrichment analysis (GSEA) software was downloaded from http://software.broadinstitute.org/gsea/. A ranking score was computed for each coding gene which had the cpm > 1 in at least 5 samples. The parameters set for each analysis were: enrichment statistic as weighted, the number of permutations was 1000, exclude sets larger than 500 and exclude sets smaller than 15. The library curated gene sets (C2) was used from Molecular Signatures Database v6.2 (MSigDB). The gene sets were statistically relevant if their FDR was below 0.05. The gene sets were considered as positively enriched if their normalized enriched score (NES) was above 1.4 and negatively enriched if their NES < -1.4 (Subramanian et al., 2005).

#### Copy number assay

DNA was isolated from bulk hESC cultures by proteinase K – SDS lysis, followed by phenol-chloroform extraction and ethanol precipitation. Copy-number quantification for *NANOG* gene was performed using TaqMan® Copy Number Assay (Hs03820140_cn). Copy number assay for RNAseP (Hs05152806_cn) was used as a reference. Samples were run on the ViiA 7 thermocycler and the results were analysed using Applied Biosystems Copy Caller v.2.1.

#### Supplemental Tables

**Table S1.** Karyotypes and passage numbers of the hESC lines used in the study.

| hESC line | Passage numbers | |
|---|---|---|
| VUB01 | Karyotyped | P70 (46, XY) |
| | Used for differentiation | P 75 - 84 |
| | Used for RNAseq (undiff) | P 75 -76 |
| | Used for RNAseq (24h ME) | P 78 - 80 |
| VUB02 | Karyotyped | P10 (46, XY) |
| | Used for differentiation | P 15 - 23 |
| | Used for RNAseq (undiff) | P 15 - 16 |
| | Used for RNAseq (24h ME) | P 17 - 23 |
| VUB03 | Karyotyped | P18 (46, XX) |
| | Used for differentiation | P 23 - 31 |
| | Used for RNAseq (undiff) | P 23 - 25 |
| VUB04 | Karyotyped | P15 (46, XX) |
| | Used for differentiation | P 17 - 28 |
| | Used for RNAseq (undiff) | P 16 - 21 |
| | Used for RNAseq (24h ME) | P 19 - 27 |

**Table S2.** Primary antibodies used in the study.

| Antibody against | Company (Cat. No.) | Concentration used |
|---|---|---|
| T (Brachyury) | R&D Systems (AF2085) | 1/200 |
| SOX2 | R&D Systems (MAB2018) | 1/100 |
| | R&D Systems (AF2018) | 1/500 |
| SOX17 | R&D Systems (AF1924) | 1/200 |
| FOXA2 | Abnova (H00003170-M12) | 1/200 |
| HNF4α | Santa Cruz (sc-374229) | 1/200 |
| β-catenin | Abcam (ab32572) | 1/250 |
| PAX6 | R&D Systems (AF8150) | 1/40 |
| POU5F1 | Santa Cruz (sc-5279) | 1/200 |

**Table S3.** TaqMan or in-house assays used in the study.

| Gene | TaqMan assay |
|---|---|
| *GUSB* | Hs99999908_m1 |
| *T* | Hs00610080_m1 |
| *SOX17* | Hs00751752_s1 |
| *FOXA2* | Hs00232764_m1 |
| *POU5F1* | Hs00742896_s1 |
| *HNF4α* | Hs00604435_m1 |
| *AFP* | Hs00173490_m1 |
| *NANOG* | Hs02387400_g1 |
| *SOX2* | Hs01053049_s1 |
| *PAX6* | Hs00240871_m1 |
| *SOX1* | Hs01057642_s1 |
| *KDR* | Hs00911700_m1 |
| *PDGFRA* | Hs00998018_m1 |
| *UBC* | In-house assay:<br>Forward 5’-CGC-AGC-CGG-GAT-TTG-3’<br>Reverse 5’-TCA-AGT-GAC-GAT-CAC-AGC-GA-3’<br>Probe 6-FAM- TCG-CAG-TTC-TTG-TTT-GTG-MGB |
